# Is Manganese the Root Cause of Human Mental Disorders?

**DOI:** 10.64898/2026.01.06.698023

**Authors:** Yaohui Bai, Xijuan Wang, Yuhan Wang, Yutong Xue

**Affiliations:** Research Center for Eco-Environmental Sciences, Chinese Academy of Sciences, Beijing, 100085, China; College of Life and Environmental Science, Minzu University of China, Beijing 100081, China; School of Eco-Environment, Harbin Institute of Technology, Shenzhen, 518055, China; National Engineering Laboratory for Advanced Municipal Wastewater Treatment and Reuse Technology, Beijing University of Technology, Beijing 100124, China

**Author notes:** Corresponding authors. (Y. Bai).

## Abstract

Mental health disorders are increasing globally and emerging as a leading public health challenge. The etiology of mental disorders is traditionally attributed to a complex interplay of genetic predispositions and social behaviors. Here, we challenge this view and propose that environmental contamination — specifically, exposure to manganese (Mn) — is a critical, yet overlooked, driver for the occurrence of mental disorders. Mn is widely used in various aspects of our lives, such as Mn steel for urbanization, agricultural fertilizer, electronic vehicle batteries, and mobile phone. Through laboratory experiments, we demonstrated that Mn exposure slightly enhanced the bacterial stress sensitivity, evidenced by altered gene expression in two-component systems and their Mn(II) oxidation function. Therefore, we hypothesized that cosmopolitan environmental Mn could increase the stress sensitivity of urban airborne microbiomes. Through comparative analysis of downloaded airborne microbiome datasets from eight cities, we found that urbanization levels show a significant positive correlation with the stress sensitivity of airborne microbiomes. Given that airborne particles can enter the human body and potentially impact health, we hypothesized that increased stress sensitivity in airborne microbiomes may influence the human gut microbiome, thereby contributing to mental disorders. To test this, we analyzed two publicly available metagenomic datasets of human gut microbiomes from both mental disorder patients and healthy controls. However, limited differences in stress sensitivity were observed between the groups, partly due to the lack of RNA-seq data for more precise functional profiling. Based on these findings, we propose a pathway linking environmental Mn exposure to increased stress sensitivity in the airborne microbiome, potentially inducing mental disorders.

## Introduction

Mental disorders — such as anxiety, depression, and bipolar disorder — represent a leading challenge to global public health, with their prevalence rising annually ^1^. More than 1 billion people worldwide live with a mental disorder ^2^. Despite their significant impact, the etiology of these disorders remains unclear. It is generally acknowledged that while genetic inheritance confers primary susceptibility, the manifestation of mental disorders is shaped by a complex interplay of individual, familial, community, and societal factors, with life experiences such as trauma acting as potential triggers ^2^. Here, we challenge this statement and propose a distinct etiology: we posit that environmental contamination, specifically the widespread use of manganese (Mn), is a fundamental cause of mental disorders.

Mn — the second most abundant transition metal in Earth’s crust —underpins core biological functions, serving in superoxide dismutase and the oxygen-evolving complex of photosystem II ^3^. Meanwhile, Mn is widely used in various aspects of our life, such as Mn steel, agricultural fertilizer ^4^, and electronic vehicle battery. Therein, Mn steel—specifically the austenitic high-Mn grade known as Hadfield steel, typically comprising 11–14% Mn—functions as a foundational material in modern urbanization and heavy industry. This alloy is distinguished by its exceptional work-hardening capability, resulting in remarkable toughness, impact resistance, and abrasion durability ^5^. As a cornerstone of modern urbanization, city dwellers are, in a sense, embedded in a vast ‘Mn reservoir’. Although previous studies in Mn mining areas have established that high-level exposure to Mn dust can cause human neurological dysfunction ^6,7^. However, the indirect health impacts of chronic exposure to Mn in these urban structures remain unreported.

Building on recent findings that environmental stress (defined as any external pressure or disturbance that negatively impacts microbial growth) sensitivity determines bacterial Mn(II) oxidation ^8^ and enhances bacterial survival ^9^, we hypothesized that the resulting Mn(IV) may act to sustain this stress sensitivity. To test this, we conducted a 154-day laboratory experiment, passaging bacteria on a series of agar media amended with chemical MnO₂ at 15-day intervals to monitor changes in bacterial stress sensitivity. Having verified the hypothesis with our results, we proposed a broader environmental impact: the extensive use of Mn in society may increase the overall stress sensitivity of airborne microbiomes by compromising their community resilience. To investigate this, we analyzed publicly available metagenomic data of airborne microbiomes from the NCBI (National Center for Biotechnology Information, USA), EMBL (European Molecular Biology Laboratory, European) and NGDC (National Genomics Data Center, China) to compare microbial stress sensitivity profiles. Finally, given the fact that the atmosphere particles can significantly influence the human gut microbiome ^10,11,12^, we subsequently compared the gut microbiomes of individuals with and without mental disorders using downloaded metagenomic datasets. We proposed a novel pathway linking environmental Mn exposure to human mental disorders via the airborne microbiome. Central to this pathway is the mechanism whereby Mn increases microbial stress sensitivity, which we supposed to be a potential underlying cause of such disorders.

## Results

### Mn presence maintaining microbial stress sensitivity

To assess the effect of MnO₂ on bacterial stress sensitivity, strains *Arthrobacter* sp. QXT-31 and *Pseudomonas* sp. QJX-1 were serially transferred over 154 days on agar plates containing either standard peptone yeast extract glucose (PYG) broth or PYG broth supplemented with MnO₂. The transferred strains were then cultured in corresponding liquid media (with or without MnCl_2_). Their biological Mn(II) oxidation activity — an indirect indicator of environmental stress sensitivity, as it is positively correlated with stress within a certain range ^8^ — was monitored monthly (Figs. S1, S2). After 140 days of cultivation, bacterial suspensions were collected for RNA-seq analysis to obtain transcripts per million (TPM) reads for bacterial two-component systems (TCS) (Figs. 1a, 1b). Finally, Mn(II) oxidation activity was measured again at the endpoint of the experiment (154 d; Figs. 1c and 1d).

**Figure 1.**
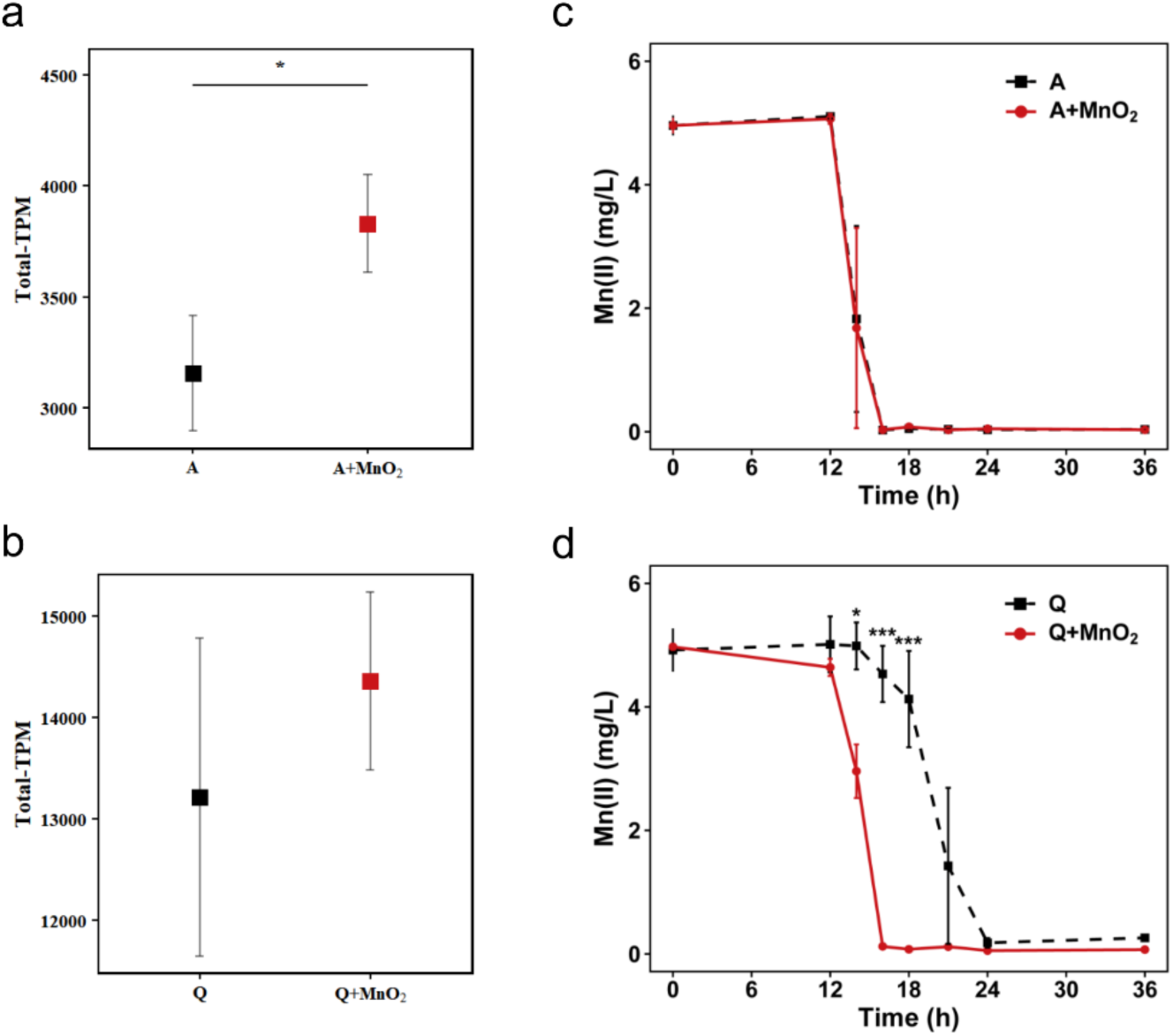
The effects of MnO_2_ presence on the stress sensitivity of bacterial strains. (a) and (b) illustrate the gene expression of two-component system (reflected in the stress sensitivity) of *Arthrobacter* sp. QXT-31 mutant (denoted as A) and *Pseudomonas* sp. QJX-1 (denoted as Q). RNA-seq was performed at experiment 140 d. (c) and (d) illustrate Mn(II) oxidation capacities of *Arthrobacter* sp. QXT-31 mutant (denoted as A) and *Pseudomonas* sp. QJX-1 (denoted as Q) at experiment 154 d. * denotes statistical significance level set at *P* < 0.05, *** denotes *P* < 0.001; T-test. Data represents the mean values of three independent biological replicates. Error bars represent the standard deviation.

The two strains exhibited distinct response patterns. For *Arthrobacter* sp. QXT-31, the total TPM (sum of all TCS transcripts) was significantly higher in the PYG+MnO₂ medium than in the PYG medium (Fig. 1a). However, its biological Mn(II) oxidation rate showed no significant difference, as Mn(II) was almost completely oxidized within 16 hours in both media (Fig. 1c). Conversely, for *Pseudomonas* sp. QJX-1, the total TPM did not differ significantly between the two media (Fig. 1b), yet its Mn(II) oxidation rate in the PYG+MnO₂ medium was significantly higher than in the PYG medium (Fig. 1d).

In summary, the addition of MnO₂ appeared to increase the environmental stress sensitivity of the bacterial strains during the experimental period, as inferred from changes in TCS gene expression and in biological Mn(II) oxidation. A more extended investigation is required to observe clearer functional differences between growth in PYG+MnO₂ and PYG media.

### Stress sensitivity of air-borne microbiome

We analyzed histidine kinase (HK) genes from two-component systems—which serve as indicators of environmental stress ^13^—in the airborne microbiomes of eight cities. After normalizing HK abundance to 16S rRNA gene levels (Fig. 2a), Guangzhou, China exhibited the lowest average HK abundance, while Hong Kong showed the highest. Notably, although the two cities are geographically close, their HK abundances differed significantly. This suggests a potential link between urban development and HK abundance in airborne microbiomes. Subsequently, we assessed the correlation between average HK abundance and urbanization level (Fig. 2b). The results revealed a statistically significant positive correlation. Furthermore, HK genes clustered into two distinct groups; however, no clear factor governing their distribution was identified (Fig. S3).

**Figure 2.**
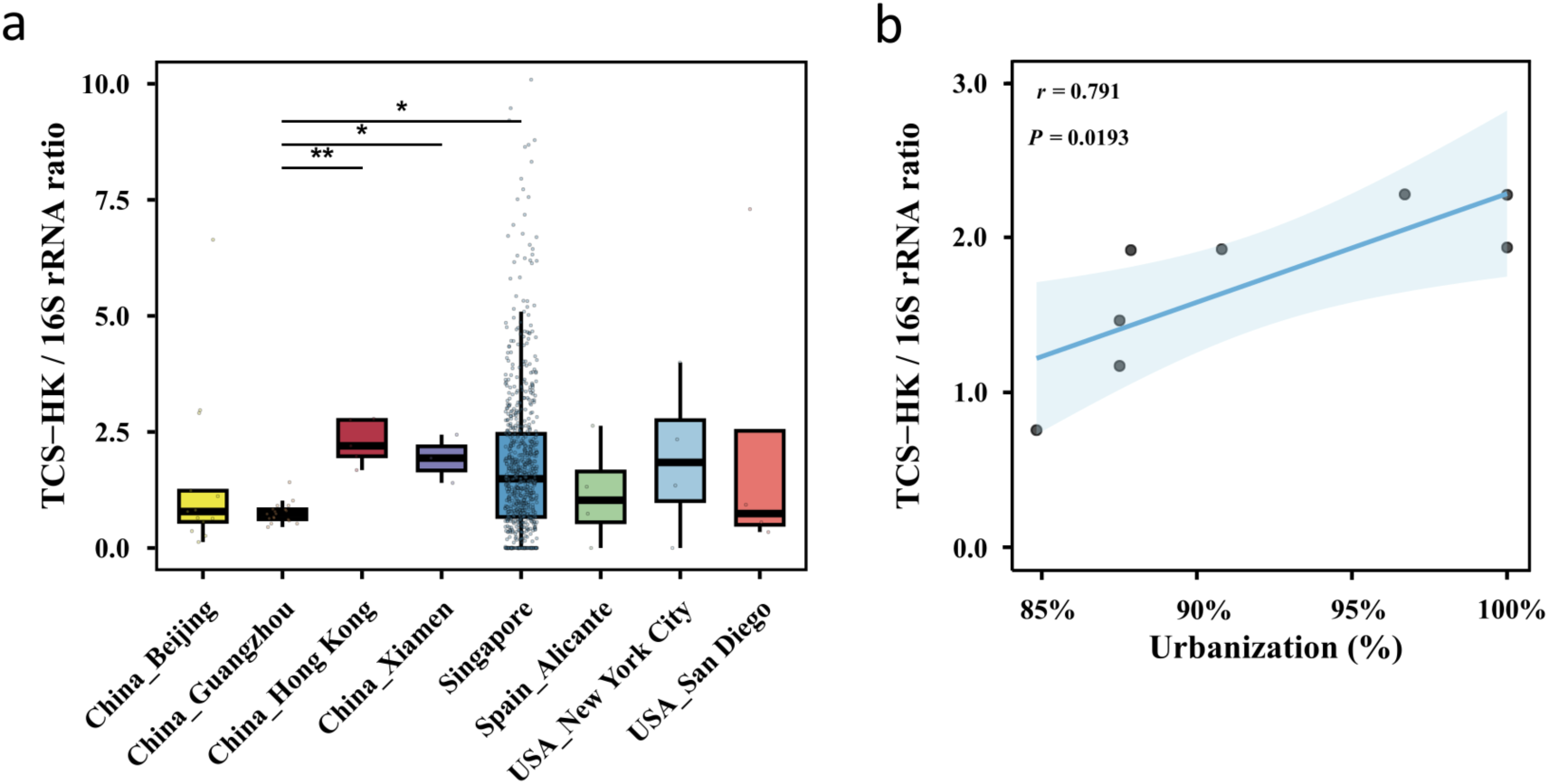
Characteristics of stress sensitivity in the air-borne microbiomes of eight cities. (a) Abundance of histidine kinase genes of two-component systems in the air-borne microbiome of eight cities. The cities and sampling time are: (1) Beijing, China (2020, 2021, 2022); (2) Guangzhou, China (2016, 2017, 2019, 2020); (3) Hong Kong, China (2016); (4) Xiamen, China (2023); (5) Singapore (2016, 2017); (6) Alicante, Spain (2016, 2017); (7) New York City, USA (2007); (8) San Diego, USA (2010). * denotes statistical significance level set at *P* < 0.05; ** denotes *P* < 0.01 using Wilcoxon–Mann–Whitney Test. (b) Correlation between the average HK gene abundance of each city and urbanization levels of each city. Details are displayed in Supporting Excel.

**Figure 3.**
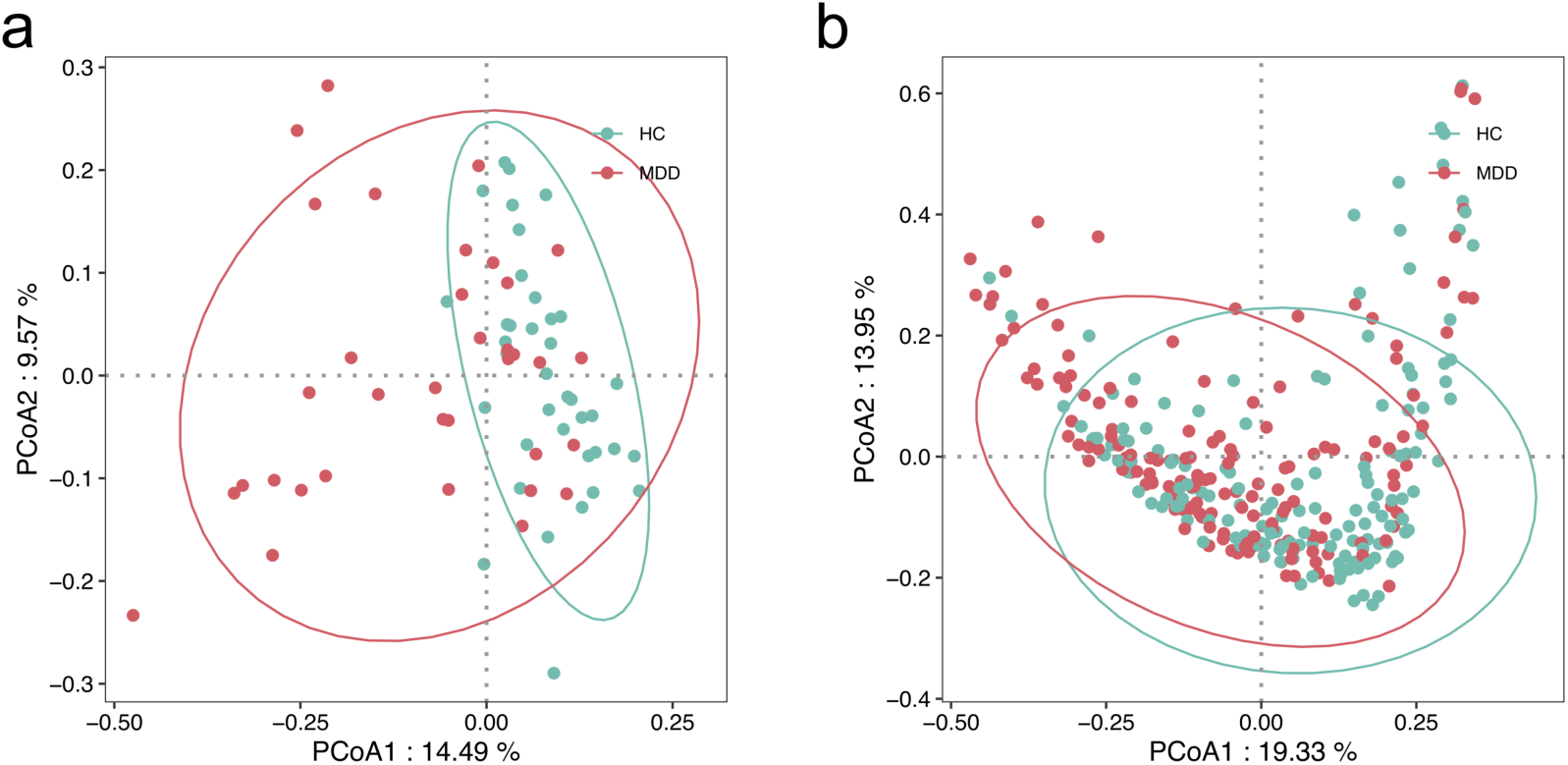
PCoA analysis of the stress sensitivity of human gut microbiome with major depression disorder and healthy controls. We downloaded two metagenomic datasets from PRJNA762199 (a) and CNP0001162 (b) and then blasted them against self-constructed two-component systems database.

### Stress sensitivity of human gut microbiome

Airborne microbes can enter the gastrointestinal tract via inhaled particles or through the consumption of food and water ^14^. This process may alter the gut microbiome. Accumulating evidence has shown that the gut microbiome is linked to mental disorders through the microbiota–gut–brain axis ^15,16,17,18,19,20,21^. In this study, we selected and downloaded two public gut metagenomic datasets associated with major depressive disorder (MDD) and health controls. Using the response regulator of two-component systems as an indicator, we assessed stress-response regulation in the gut microbiome between individuals with MDD and healthy controls (HCs). Analysis of the PRJNA762199 dataset revealed a partial separation between some MDD samples and HCs (PERMANOVA p = 0.001, R² = 0.06). In contrast, no clear separation was observed in the CNP0001162 dataset, where HCs clustered together with MDD samples (PERMANOVA p = 0.386, R² = 0). This discrepancy may be attributed to the absence of RNA-seq data in our analysis, as metagenomic data reflect genetic potential (DNA) rather than actual gene activity. Therefore, future studies incorporating RNA-seq are warranted to elucidate expression-level differences between MDD and HC groups.

## Discussion

### Pathway inducing human mental disorder

Accumulating evidence indicates that functional alterations in the gut microbiome are a key driver in the development of mental disorders ^19,22,23^. However, the question of “how and which factors induce these changes in the human gut microbiome” remains unresolved. Here, we propose that airborne microbes could be a pivotal factor. These microbes can enter the human body and subsequently influence the structure and function of the microbiome in various organs ^24^. Given the extensive use of Mn in modern life, the overall environmental stress on airborne microbial communities is gradually increasing, as Mn may compromise their community stability and resilience. This, in turn, could heighten the environmental stress sensitivity of the gut microbiome and undermine its resilience—the ability to recover from disturbance.

Since gut microbiome resilience significantly impacts psychological symptoms, emotion regulation, and cognitive function ^25^, its disruption could indeed contribute to mental disorders. In addition, airborne microbes might directly affect the central nervous system. The possibility that these microbes could enter the brain raises an intriguing question deserving of further discussion: whether a resident brain microbiome exists ^26,27^.

Overall, mental disorders likely result from the combined effects of both the direct impact of these microbes and their indirect influence via the microbiota–gut–brain axis. Therefore, we propose a putative pathway: Mn exposure → increased environmental stress on the airborne microbiome → elevated stress on the gut (and potentially brain) microbiome → increased susceptibility to mental disorders (Fig. 4). This proposed pathway requires further validation; thus, large-scale cohort studies are warranted.

**Figure 4.**
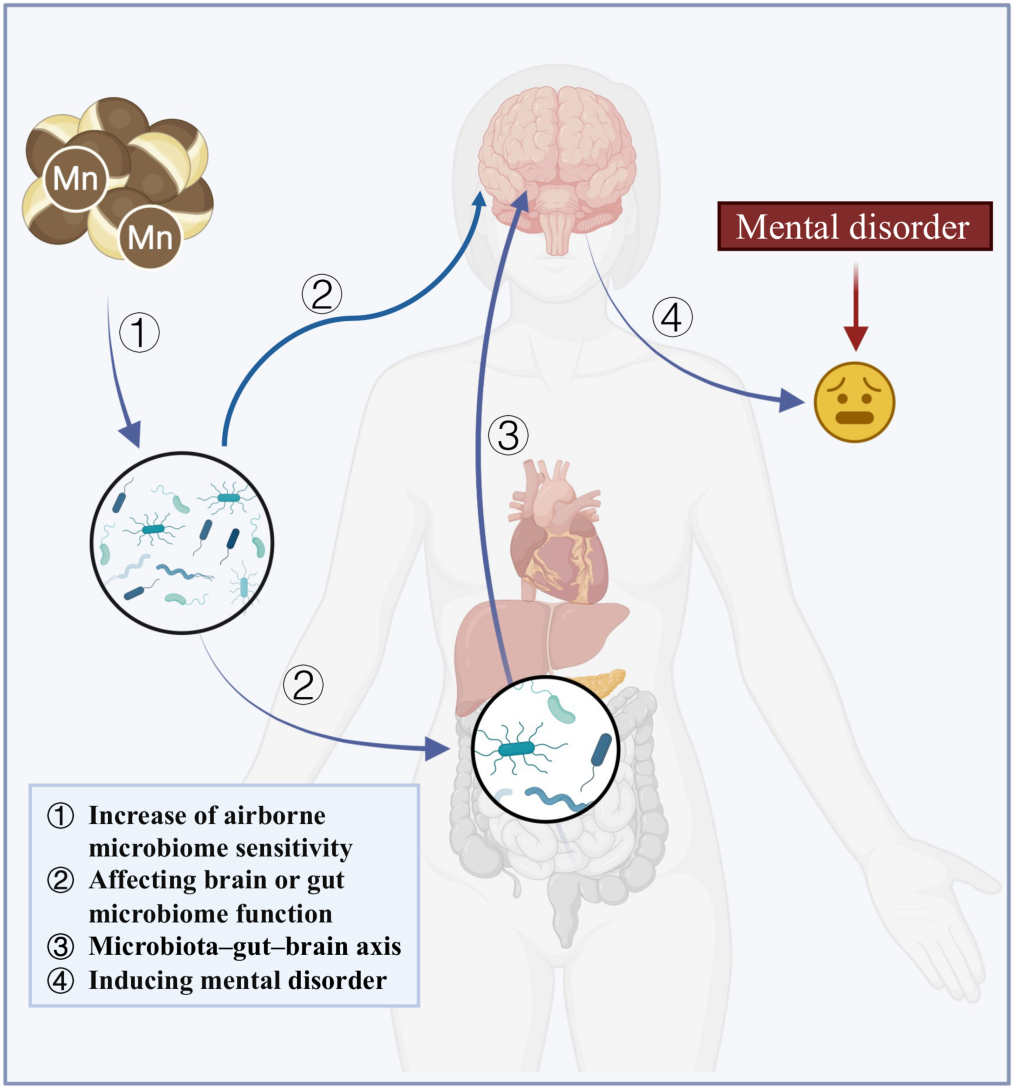
Proposed “Mn exposure → increasing environmental stress of airborne microbiome → increasing environmental stress of gut microbiome or brain microbiome → inducing mental disorder” pathway.

### Urbanization heightening stress sensitivity

Urbanization is accelerating globally, driven by its potential to stimulate economic growth. Many countries, including the United States, China, and South Korea, have actively promoted urbanization as a core development strategy. However, evidence suggests that urbanization and the accompanying economic growth do not automatically lead to sustained improvements in overall well-being ^28^. Extensive use of materials such as Mn steel in urban construction may alter the ambient airborne microbiome, potentially increase environmental stress sensitivity and contribute to mental health disorders. Despite these concerns, many rapidly urbanizing countries in Africa and elsewhere view it as essential for enhancing national strength. We propose that an optimal level of urbanization likely exists — one that balances economic development with the well-being of residents. Finding this equilibrium is crucial for achieving long-term, sustainable progress.

### Heightened stress sensitivity inducing abnormal human behavior

Heightened stress sensitivity can induce significant behavioral changes. Stress sensitivity exerts a bipolar influence: on one hand, it may enhance cognitive alertness, potentially linked to longer lifespans; on the other, it can lead to increased aggression and abnormal behaviors, such as too sensitive to stress. Currently, young adults worldwide are experiencing a rising tide of mental health issues ^29^. A diminished willingness to have children is observed in this demographic. We postulate that the presence of Mn, by creating overwhelming stress, is a primary driver of this phenomenon. If this is the case, purely material incentives would be ineffective in reversing the trend.

### Limitation

Our proposed pathway has not been fully confirmed, primarily because of the limited availability of public RNA-based microbiome data. In particular, the current evidence is limited to observational cues indicating that stress sensitivity in both airborne and gut microbial communities tends to increase over time.

In summary, we challenge the traditional etiological view of mental disorders, which emphasizes a complex interplay between genetic predisposition and social behavior. Rather, we propose that environmental contamination — specifically, exposure to Mn — constitutes a fundamental causative factor. However, the lack of publicly available temporal RNA-seq data on both the airborne and human gut microbiomes means this pathway requires further confirmation.

We advocate enhanced sharing of multi-omics data, particularly RNA-sequencing data from both atmospheric and human-gut microbiomes, to advance the etiological research of mental disorders. Also, we urge scholars across nations and disciplines to unite by setting aside academic biases, to tackle the root causes of mental disorders. Time is of the essence; we must take immediate action to protect lives.

## Materials and Methods

### Culture medium and bacterial strains

*Pseudomonas* sp. QJX-1 ^30^ and *Arthrobacter* sp. QXT-31 mutant ^31^ strains used in this study were detailed described in our previous studies. These two bacterial strains encode the Mn(II)-oxidizing genes and can induce Mn(II) oxidation activity individually. Both strains were initially cultured in the modified PYG medium ^32^ (500 mg/L of MgSO_4_·7H_2_O, 60 mg/L of CaCl_2_·2H_2_O and 0.25 g/L of peptone, yeast extract and glucose) buffered by 10 mM 4-(2-hydroxyethyl)-1-piperazineethanesulfonic acid (HEPES) (pH 7.2). All the cultivation experiments were performed in the dark shakers.

### Impact of Mn presence on stress sensitivity of bacterial strains

The experiment was conducted using PYG media, both with and without Mn (Mn) supplementation. To prepare the Mn-supplemented medium, sterile PYG agar (1.8%) was first poured into Petri dishes, after which 0.005 M of solid MnO₂ was evenly applied to the surface (Fig. S4). We then overlaid the solid MnO₂ with a layer of agar (∼0.2 cm). Two bacterial strains, *Pseudomonas* sp. QJX-1 and *Arthrobacter* sp. QXT-31 mutant, were initially cultured overnight in 150-mL flasks. The bacterial suspensions were then streaked onto fresh PYG agar plates, either with or without MnO₂. These strains were subsequently transferred to fresh corresponding agar plates at about 15-day intervals (Fig. S5). After each monthly transfer, individual colonies were selected from the agar plates and inoculated into liquid PYG media, with or without MnCl_2_. for further analysis. The bacterial stress sensitivity was assessed indirectly by measuring Mn(II) oxidation at 24, 48, and 72 hours, based on our previous finding of a positive correlation between Mn(II) oxidation and stress sensitivity ^8^. To further validate the increase in sensitivity, RNA was extracted from the bacterial cultures at the 72-hour time point. Specifically, cell pellets were harvested by centrifugation at 10,000 × *g* for 5 minutes at 4°C and immediately processed for RNA extraction using TRNzol Reagent (TIANGEN, Beijing, China), following the manufacturer’s instructions.

### RNA-seq

RNA integrity was evaluated using an Agilent 2100 Bioanalyzer (Agilent Technologies, Santa Clara, CA, USA), and only samples with RNA integrity number (RIN) or RNA quality number (RQN) values greater than 8.0 were selected for cDNA library construction and sequencing. Libraries were sequenced on the Illumina NovaSeq 6000 platform, generating 250-bp paired-end reads. Raw sequence reads were processed to remove adapter sequences and low-quality reads using fastp ^33^.

### Quantifying the expression levels for two-component system genes from transcriptome data of two bacterial strains

To obtain the gene sequences necessary for functional identification, gene prediction for the reference genome was first performed de novo using prodigal ^34^ (v2.6.3) to generate GFF3 annotation files, along with coding sequences. To identify specific functional genes from the reference genome, the coding sequences were aligned against a specialized prokaryotic two-component system database -- P2CS database ^35^ using blastn (v2.15.0) with an E-value cutoff of 1e-20, thereby identifying genes associated with two-component systems.

Subsequently, clean sequencing reads of RNA-seq data were aligned to the genome using bowtie2 ^36^ (v2.5.3). The alignment results were processed with SAMtools ^37^ (v1.20), and raw counts for each gene were quantified using featureCounts ^38^ (v2.1.1). Gene expression levels were normalized using the Transcripts Per Million (TPM) measure based on the raw counts. The TPM for each gene was calculated using Equation 2:

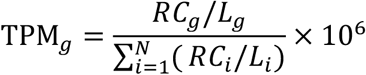

Where RC_g_ is the raw read count mapped to gene g, L_g_ is the length of gene g in base pairs, and N is the total number of genes. This two-step calculation normalizes read counts by gene length (yielding Reads Per Kilobase, RPK), and then scales the sum of all RPK values in the sample to one million.

Finally, based on the identified target gene identifiers, the expression levels of two-component system genes were extracted and aggregated from the genome-wide TPM matrix for downstream analysis.

### Quantification of two-component system gene abundance in metagenomic data

Metagenomic reads were aligned against P2CS database ^35^, using blastn ^39^ (v2.15.0) with a recommended e-value cutoff of 1e-20. Alignments were further filtered to retain only those with >70% identity. The resulting TCS gene abundance was calculated using the following Equation:

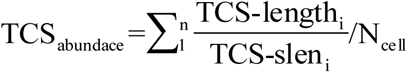

Where TCS-slen_i_ represents length of matched reference sequence in the database, and TCS-length_i_ represents sum of alignment length of all input sequences annotated as the reference gene, N_cell_ represents the estimation results of cell count output by MnOxGeneTool ^40^.

### Stress sensitivity of air-borne microbiome

Given the lack of publicly available RNA-seq data, our analysis was restricted to comparing histidine kinases of two-component systems (TCSs) – which are indicative of environmental stress response – from the metagenomic datasets of eight cities. We systematically identified 834 metagenomic samples from 7 studies on outdoor airborne bacterial communities by searching published literature and the NCBI database with keywords including "air", "airborne", and "atmosphere". These samples originate from outdoor air environments in 8 cities worldwide. Following retrieval from the NCBI SRA, raw sequencing data were converted to FASTQ format using fasterq-dump (-split-3) and processed for quality control (including removal of low-quality sequences and adapter contamination) using fastp (v0.23.1). TCS-related genes were then quantified using the above method. By retrieving publicly available data from relevant statistical yearbooks and population censuses, quantitative indicators of urbanization levels for the sampled cities corresponding to the study samples in the sampling years were extracted (here, the proportion of urban population was selected as the indicator). Pearson correlation analysis was performed to examine the linear relationship between the level of urbanization and the gene abundance of histidine kinase (HK) in the bacterial TCS. Using the cor.test function in R (version 4.5.2), the correlation coefficient, statistical significance (*p*-value), and 95% confidence interval were calculated.

### Stress sensitivity of gut microbiome in individuals with and without mental disorders

To investigate the stress sensitivity of the gut microbiome in individuals with mental disorders, we obtained two metagenomic datasets: PRJNA762199 from the NCBI and CNP0001162 from the China National GeneBank (CNGB). The first dataset (PRJNA762199) comprises 36 individuals with major depressive disorder (MDD) and 36 healthy controls (HC). Notably, none of the patients had undergone medication prior to fecal sample collection ^41^. The second dataset (CNP0001162) includes 154 MDD patients and 151 HC. In this cohort, all patients were unmedicated, and no significant differences were observed in demographic indices—such as gender, age, or body mass index—between the patient and control groups ^42^. We analyzed their TCS expression levels from the metagenomic reads, as described above.

### Statistical analysis

All statistical analyses and visualizations were conducted in the R 4.5.2. The T-test was used for assessing of environmental stress sensitivity and biological Mn(II) oxidation of two bacterial strains, with the statistical significance level set at *p* < 0.05. The Wilcoxon–Mann–Whitney Test was used for assessing environmental stress sensitivity in different cities, with the statistical significance level set at *p* < 0.05.

## Supporting information

Supplemental Word

## Data availability

All raw reads were deposited in the China National Center for Bioinformation database (https://www.cncb.ac.cn) under BioProject PRJCA052540. Transcriptome sequence data for *Arthrobacter* sp. QXT-31 mutant and *Pseudomonas* sp. QJX-1, have been deposited in under accession number CRX2306118 - CRX2306113 and CRX2306124 - CRX2306119, respectively. Unique biological materials are available from the corresponding author upon reasonable request. [Source data](https://www.nature.com/articles/s41586-020-2468-5#Sec26) are provided with this paper.

## Acknowledgments

This work was supported by the National Natural Science Foundation of China (52450009). We kindly thank Linhao Zhang for his valuable helps in bioinformatics analysis.

## Author contributions

Y.B. conceived and designed the study. Y.X., Y.B., Y.W., and X.W. carried out experiments on the impact of Mn on bacterial stress sensitivity and performed corresponding data analysis. X.W., Y.B., and Y.W. retrieved public metagenomic data from airborne microbiome samples and evaluated microbial stress sensitivity. Y.B., and Y.W. retrieved public metagenomic data from human gut microbiome samples and evaluated microbial stress sensitivity. The manuscript was drafted by Y.B. All authors read, reviewed, and approved the final version of the manuscript.

## Competing interests

The authors declare no competing interests.

## Notes

During the preparation of this work, the authors used AI Deepseek to improve readability and language. After using this tool, the authors reviewed and edited the content as needed and take full responsibility for the content of the publication.

## Notes

### Competing Interest Statement

The authors have declared no competing interest.

## References

1 Vos, T. et al. Global burden of 369 diseases and injuries in 204 countries and territories, 1990&#x2013;2019: a systematic analysis for the Global Burden of Disease Study 2019. The Lancet 396, 1204–1222 (2020).

2. Organization;, W. H. (Geneva, 2025).

3 Tebo, B. M., Geszvain, K. & Lee, S.-W. The molecular geomicrobiology of bacterial manganese (II) oxidation. (Springer, 2010).

4 Rezende, V. P. et al. Reviewing Manganese and Zinc Foliar Fertilization Approaches and Results through a Quantitative Metadata Analysis. ACS Agricultural Science & Technology 5, 1792–1802 (2025).

5 Jacob, R., Raman Sankaranarayanan, S. & Kumaresh Babu, S. P. Recent advancements in manganese steels – A review. Materials Today: Proceedings 27, 2852–2858 (2020).

6 Harischandra, D. S. et al. Manganese-Induced Neurotoxicity: New Insights Into the Triad of Protein Misfolding, Mitochondrial Impairment, and Neuroinflammation. Front Neurosci-Switz 13 (2019).

7 Miah, M. R. et al. The effects of manganese overexposure on brain health. Neurochem Int 135, 104688 (2020).

8 Bai, Y. et al. Environmental stress sensitivity determines bacterial Mn (II) oxidation. 2025.2005. 2009.653071 (2025).

9 Lin, H. & Bai, Y. Physiological significance of bacterial Mn(II) oxidation. bioRxiv, 2025.2011.2024.690301 (2025).

10 Fouladi, F. et al. Air pollution exposure is associated with the gut microbiome as revealed by shotgun metagenomic sequencing. Environment International 138, 105604 (2020).

11 Dujardin, C. E. et al. Impact of air quality on the gastrointestinal microbiome: A review. Environmental Research 186, 109485 (2020).

12 Salim, S. Y., Kaplan, G. G. & Madsen, K. L. Air pollution effects on the gut microbiota: a link between exposure and inflammatory disease. Gut Microbes 5, 215–219 (2014).

13. Kaserer, A. O. & West, A. H. in Handbook of Cell Signaling (Second Edition) (eds Ralph A. Bradshaw & Edward A. Dennis) 581–586 (Academic Press, 2010).

14 Albano, G. D., Montalbano, A. M., Gagliardo, R., Anzalone, G. & Profita, M. Impact of Air Pollution in Airway Diseases: Role of the Epithelial Cells (Cell Models and Biomarkers). Int J Mol Sci 23 (2022).

15 Andrioaie, I. M. et al. The Role of the Gut Microbiome in Psychiatric Disorders. Microorganisms 10 (2022).

16 Bear, T. et al. The Microbiome-Gut-Brain Axis and Resilience to Developing Anxiety or Depression under Stress. Microorganisms 9 (2021).

17 Xiong, R. G. et al. The Role of Gut Microbiota in Anxiety, Depression, and Other Mental Disorders as Well as the Protective Effects of Dietary Components. Nutrients 15 (2023).

18 Afroz, K. F. & Manchia, M. Gut microbiome and psychiatric disorders. Bmc Psychiatry 23, 488 (2023).

19 Kamath, S. et al. Distinguishing the causative, correlative and bidirectional roles of the gut microbiota in mental health. Nature Mental Health 3, 1137–1151 (2025).

20 Pennisi, E. Gut bacteria linked to mental well-being and depression. Science 363, 569–569 (2019).

21 Ohara, T. E. & Hsiao, E. Y. Microbiota–neuroepithelial signalling across the gut–brain axis. Nature Reviews Microbiology 23, 371–384 (2025).

22 Kumar, A. et al. Gut Microbiota in Anxiety and Depression: Unveiling the Relationships and Management Options. Pharmaceuticals (Basel) 16 (2023).

23 Ma, Z., Zhao, H., Zhao, M., Zhang, J. & Qu, N. Gut microbiotas, inflammatory factors, and mental-behavioral disorders: A mendelian randomization study. Journal of Affective Disorders 371, 113–123 (2025).

24 Costa, L. G. et al. Effects of air pollution on the nervous system and its possible role in neurodevelopmental and neurodegenerative disorders. Pharmacol Ther 210, 107523 (2020).

25 An, E. et al. Stress-resilience impacts psychological wellbeing as evidenced by brain–gut microbiome interactions. Nature Mental Health 2, 935–950 (2024).

26 Link, C. D. Is There a Brain Microbiome? Neurosci Insights 16, 26331055211018709 (2021).

27 The Lancet, M. Brain microbiome: is it all in our heads? The Lancet Microbe 6, 101075 (2025).

28 Sarracino, F. & J. O’Connor, K. Governments should prioritize well-being over economic growth. Nature Human Behaviour 9, 2003–2005 (2025).

29 Pei, R. et al. Bridging the empathy perception gap fosters social connection. Nature Human Behaviour 9, 2121–2134 (2025).

30 Chang, Y., Bai, Y., Huo, Y. & Qu, J. Benzophenone-4 promotes the growth of a *Pseudomonas* sp. and biogenic oxidation of Mn(II). Environmental Science & Technology 52, 1262–1269 (2018).

31 Bai, Y. et al. Environmental stress sensitivity determines bacterial Mn(II) oxidation. bioRxiv, 2025.2005.2009.653071 (2025).

32 Liang, J., Bai, Y., Men, Y. & Qu, J. Microbe–microbe interactions trigger Mn(II)-oxidizing gene expression. The ISME Journal 11, 67–77 (2017).

33. Chen, S. Ultrafast one-pass FASTQ data preprocessing, quality control, and deduplication using fastp. iMeta 2, e107 (2023).

34 Hyatt, D. et al. Prodigal: prokaryotic gene recognition and translation initiation site identification. BMC bioinformatics 11, 119 (2010).

35 Ortet, P., Whitworth, D. E., Santaella, C., Achouak, W. & Barakat, M. P2CS: updates of the prokaryotic two-component systems database. Nucleic Acids Research 43, D536–D541 (2015).

36 Langmead, B. & Salzberg, S. L. Fast gapped-read alignment with Bowtie 2. Nature methods 9, 357–359 (2012).

37 Li, H. et al. The sequence alignment/map format and SAMtools. bioinformatics 25, 2078–2079 (2009).

38 Liao, Y., Smyth, G. K. & Shi, W. featureCounts: an efficient general purpose program for assigning sequence reads to genomic features. Bioinformatics 30, 923–930 (2014).

39 Altschul, S. F., Gish, W., Miller, W., Myers, E. W. & Lipman, D. J. Basic local alignment search tool. Journal of molecular biology 215, 403–410 (1990).

40 Wang, Y. et al. MnOxGeneTool: A comprehensive tool for identifying and quantifying Mn (II)-oxidizing genes, revealing phylogenetic diversity and environmental drivers of Mn (II)-oxidizers. Environmental Science & Technology

41 Kovtun, A. S. et al. Alterations of the composition and neurometabolic profile of human gut microbiota in major depressive disorder. Biomedicines 10 (2022).

42 Yang, J. et al. Landscapes of bacterial and metabolic signatures and their interaction in major depressive disorders. Science Advances 6, eaba8555 (2020).

