## Supplemental Word for "Is Manganese the Root Cause of Human Mental Disorders?"

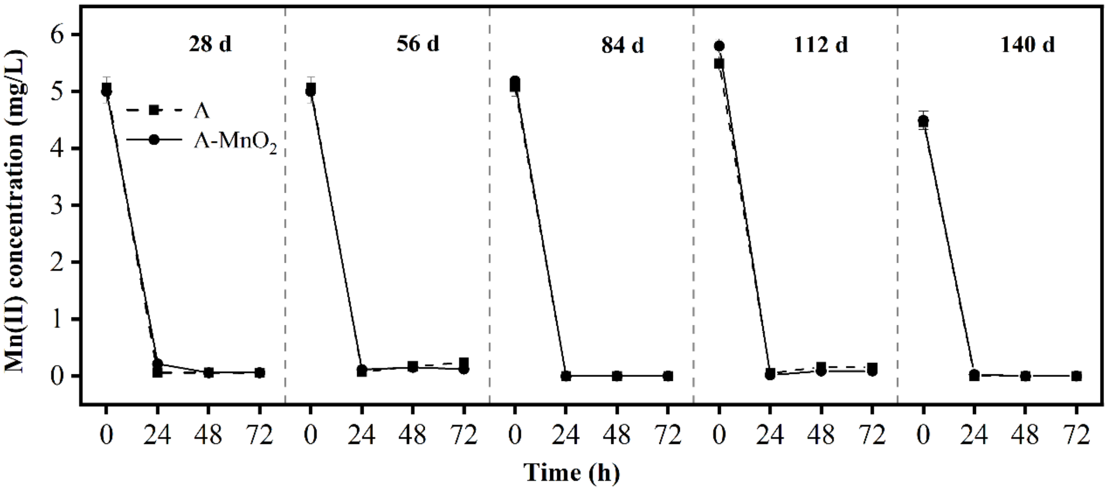


**(a)**

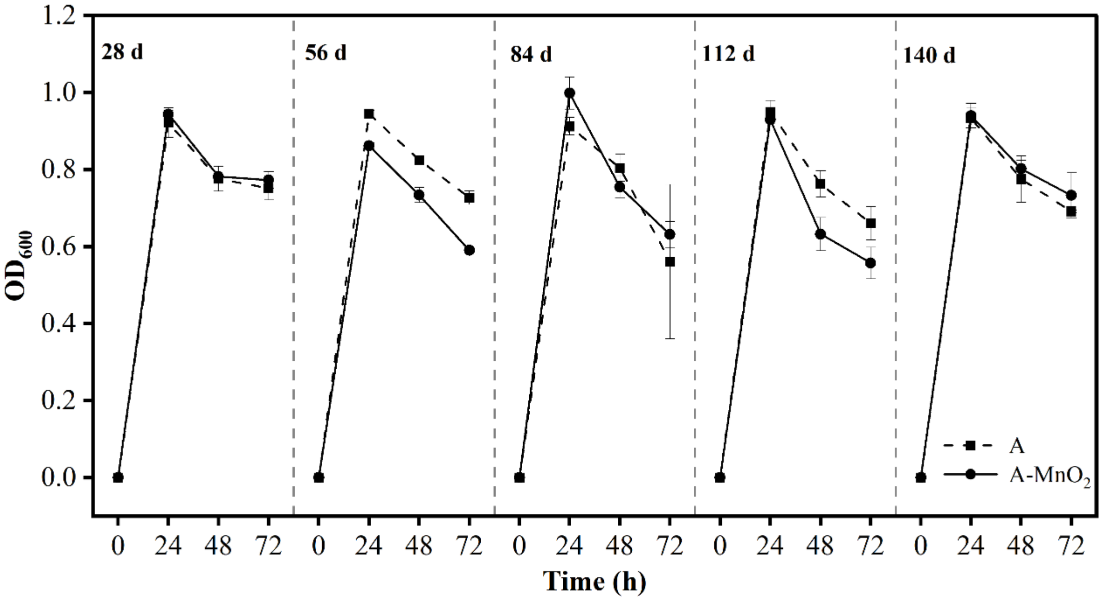


**(b)**

**Figure S1.** Mn(II) oxidation (a) and growth (b) of *Arthrobacter* sp. QXT-31 mutant (denoted as A) over time. Data represents the mean values of three independent biological replicates. Error bars represent the standard deviation.

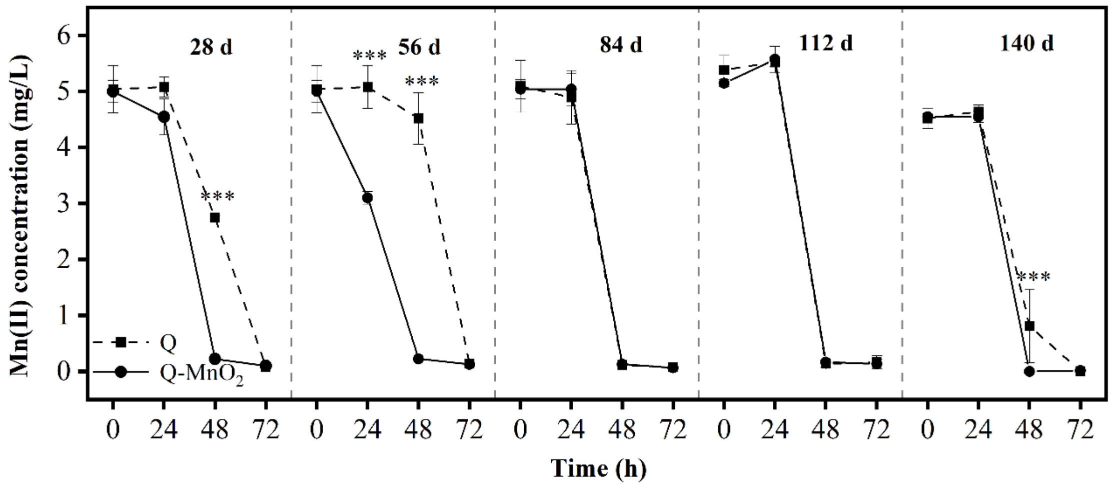


(a)

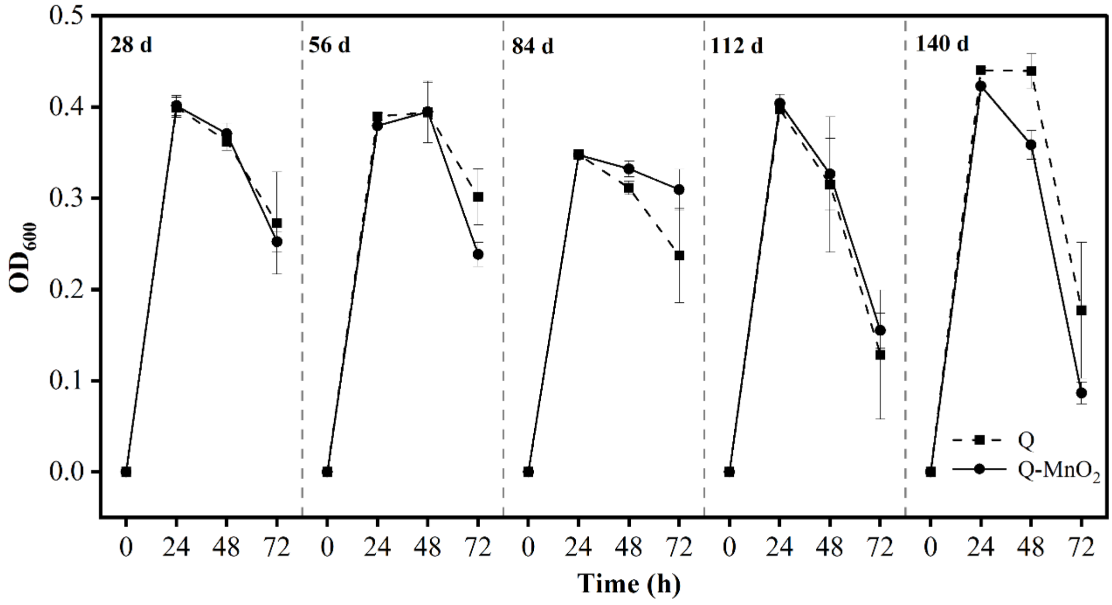


(b)

**Figure S2.** Mn(II) oxidation (a) and growth (b) of *Pseudomonas* sp. QJX-1 (denoted as A) over time. Three asterisks (***) indicate $p < 0.001$, T-test. Data represents the mean values of three independent biological replicates. Error bars represent the standard deviation.

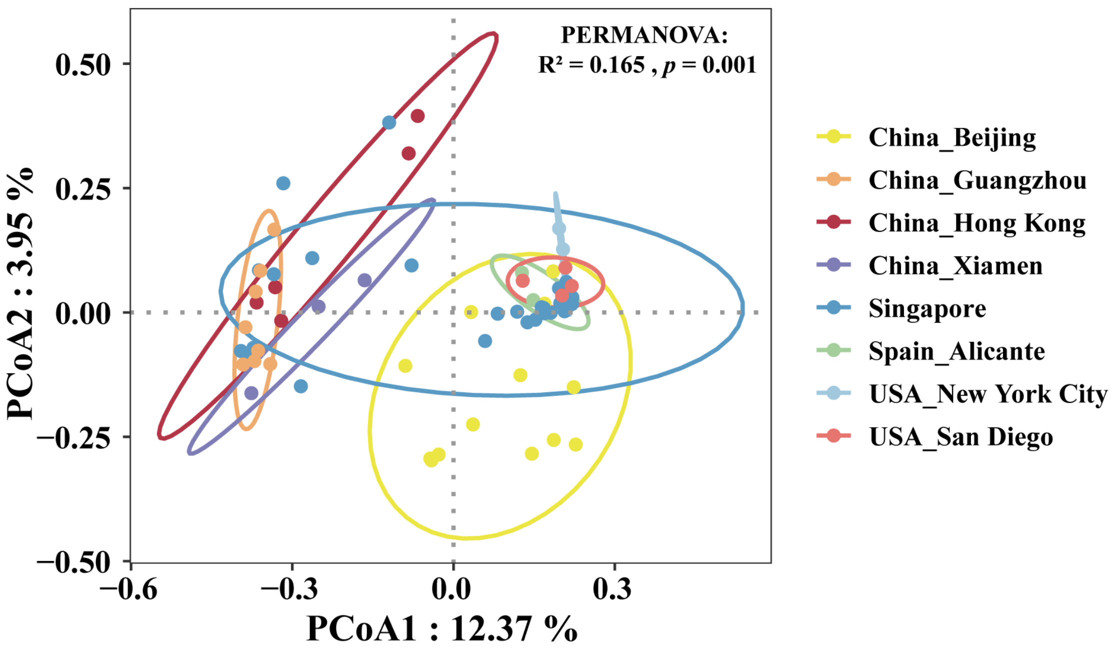


**Figure S3.** Compositional differences in Histidine Kinase (HK) Genes within two-component systems (TCS) of airborne bacteria. The sampling sites and sampling years are as follows: (1) Beijing, China (2020, 2021, 2022); (2) Guangzhou, China (2016, 2017, 2019, 2020); (3) Hong Kong, China (2016); (4) Xiamen, China (2023); (5) Singapore (2016, 2017); (6) Alicante, Spain (2016, 2017); (7) New York City, USA (2007); (8) San Diego, USA (2010).

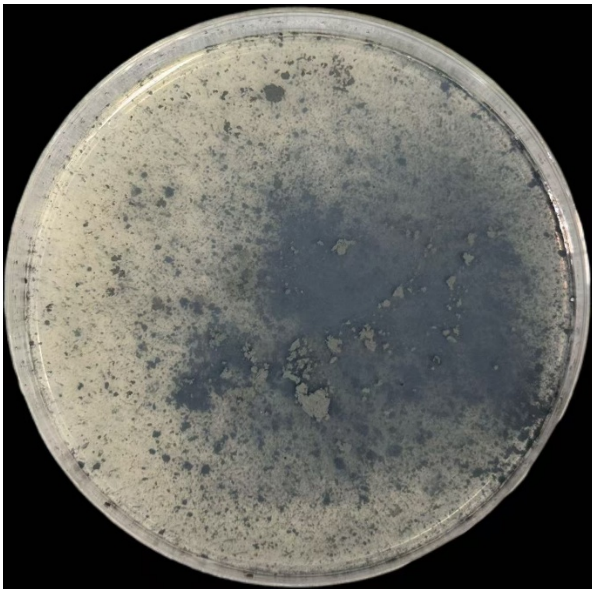

**Figure S4.** Photograph depicting the addition of $MnO_2$ onto an agar plate.

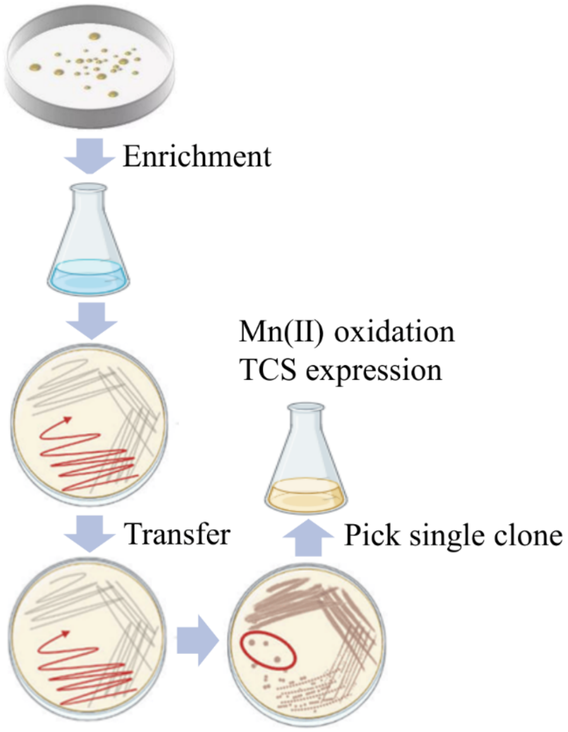


**Figure S5.** Culture experiment protocol
